## Supplementary information for "Multi-Scale Modeling of Intensive Macroalgae Cultivation and Marine Nitrogen Sequestration"

Meiron Zollmanna*, Boris Rubinskyb, Alexander Liberzonc and Alexander Golberga.

Corresponding author: Meiron Zollmann


### Supplementary Information Text

#### Methods

**Model dimensions.** The multi-scale model is a 3D model, as described in **Fig. S1**. x axis is the direction of the flow, along which cultivation reactors are chained, y axis is the horizontal direction perpendicular to the flow, along which multiple rows of reactors can be placed, and z axis points downwards from the water surface. Only one layer of reactors can be placed along the z axis, as sun exposure is necessary.

**Step-by-step multi-scale model formulation.**
*Ulva metabolism model*. Our multi-scale model was developed based on a mathematical model of metabolism of the *Ulva* sp. macroalgae, modified from previous works[1]. The model assumes that the dynamics of the limiting nutrient, in this case nitrogen (N), under the constraining effects of major environmental conditions (light intensity (I), temperature (T) and salinity (S)) predicates the dynamics of biomass growth and chemical composition. The model follows the Droop Equation concept, relating growth to internal nutrient concentrations ("cell quota") rather than to the external, environmental, concentrations[2,3]. A schematic description of this metabolism model is presented in **Fig. S2.** The model is based on three governing equations, describing the mass balance of three state variables: biomass density (m, g Dry Weight (DW) $l^{-1}$, eq S1), biomass internal nitrogen ($N_{\mathrm{int}}$, $\%\ \mathrm{g\ N\ kg\ DW^{-1}}$, eq S2) and external nitrogen in water ($N_{\mathrm{ext}}$, $\mathrm{\mu mol\ N}\ l^{-1}$, eq S3), under varying temperatures, light intensities and salinities. The set of governing ordinary differential equations (ODEs) is summarized in **Table S1.** Further details and reasoning are presented below**.**
*Growth Rate*. The specific growth rate of the algae is a function of I, T, S and nutrients [1]. In the marine environment, the limiting nutrient is usually N[4] and our model focuses on N limited environments. However, similar models can be developed also for other elements such as phosphorus (P) and ferrous (Fe) that may limit growth in some marine environments. Following Liebig's law of the minimum, we formulate the growth rate as a function of the limiting factor, which is the scarcest resource in relation to the macroalgae's requirement. This formulation is based on a simplifying assumption that there are no interactions between these factors and thus no co-limitations. However, temperature and salinity effects were excluded from the minimum law, as they constitute background conditions for metabolic or uptake efficiency and are not considered as growth resources. Therefore, the approximated function for biomass growth rate appears in eq S1.1. We adjusted daily specific growth and losses rates to hourly rates, assuming for simplicity that growth and biomass losses occur only during light hours (see details below). This assumption ignores night growth that occurs due to metabolites produced during light-time photosynthesis[5], and thus distorts growth distribution along the day. However, it has no effect on total daily growth and therefore does not impair the model accuracy in resolutions of days to weeks.

$$\mu = \mu_{max} f_{Temp} f_S \min\{f_{N_{int}}, f_{P_{int}}, f_I\} \qquad \text{(S1.1)}$$

Where $\mu_{max}$ ($h^{-1}$) is the maximum specific growth rate and $f_{Temp}$, $f_S$, $f_{N_{int}}$, $f_{P_{int}}$ and $f_I$ are the temperature, salinity, internal N, internal P and light intensity growth functions.

*Biomass Losses.* The specific rate of biomass losses due to respiration, exudation and mortality is a function of temperature, and is calculated relatively to a known losses rate in a reference temperature. We used the losses function from Martins and Marques [6]. This does not include losses by grazing, sporulation events and fragmentation by storms, which vary significantly between different environments and are highly affected by extreme events.

$$\lambda = \lambda_{20}\, \theta^{T-20} \tag{S1.2}$$

Where $\lambda_{20}$ ($h^{-1}$) is the specific rate of biomass losses and $\theta$ is an empiric factor of biomass losses.

*Temperature Effects.* Temperature affects all metabolic processes and thus also growth rate. We adapted the temperature function from Martins and Marques[6] with minor adjustments, setting the exponent $n$ as a free parameter, as opposed to the original $n = 2$.

$$f_{Temp} = \exp\left(-2.3\left(\frac{T - T_{opt}}{T_x - T_{opt}}\right)^n\right) \tag{S1.3}$$

Where $T_x = T_{min}$ for $T \le T_{opt}$ and $T_x = T_{max}$ for $T > T_{opt}$. $T_{min}$, $T_{opt}$ and $T_{max}$ (℃) are the minimal, optimal and maximal temperatures for *Ulva* growth.

*Salinity Effects.* Similarly, salinity has a general effect on nutrient uptake and growth of macroalgae. However, the effect of salinity on *Ulva* is relatively minor, as its flexible cell membrane allows it to adjust to a wide range of salinities[7]. Here, we used the salinity growth function from Martins and Marques[6] and adjusted it to fit to high salinities by changing the exponent of the original function (from $b = 2$ to $b = 4.4$) to achieve a minimal error compared to measured data from Choi et al.[7]

$$f_S = 1 - \left(\frac{S - S_{opt}}{S_x - S_{opt}}\right)^b \; for\, S \ge 5 \; or \; \frac{S - S_{min}}{S_{opt} - S_{min}} \; for\, S < 5 \tag{S1.4}$$

Where $S_x = S_{min}$ and $b = 2.5$ for $S < S_{opt}$, and $S_x = S_{max}$ and $b = 4.4$ for $S \ge S_{opt}$. $S_{min}$, $S_{opt}$ and $S_{max}$ ($PSU$) are the minimal, optimal and maximal salinities for *Ulva* growth.

*Light Effects.* Light intensity effect on growth is described by eq S1.5[8]. A more precise description of light effects will relate also to the photosynthesis process, that occurs only during light time but can result in production of new biomass also at night. For simplicity, we formulate a direct effect of light on growth, and normalized daily growth rate to per light-hour growth rate, thus minimizing the effect on model results (See growth rate section).

$$f(I) = \frac{I}{K_I + I} \tag{S1.5}$$

Where $I$ is the incident light intensity and $K_I$ (μmol photons $m^{-2}$ $s^{-1}$) is the light half saturation constant.

*Nutrients Effects.* The model assumes that at each time point growth is limited by a single factor, which is a single nutrient or the light intensity. To simplify the model, we assume that both ambient and internal N:P ratio are smaller then 12, and therefore N is the limiting nutrient and $f_{P_{int}}$=1 [1]. Internal N and light intensity effects on growth are described in eq S1.6[9]. Internal N effects can be divided into three ranges: $N_{\text{int}} = N_{\text{int min}}$ where there is no growth as reserves are empty and all $N_{\text{int}}$ is structural N, $N_{\text{int min}} < N_{\text{int}} < N_{\text{int crit}}$ where growth rate increases as $N_{\text{int}}$ increases, and $N_{\text{int crit}} < N_{\text{int}}$ where further filling of the reserves does not increase growth rate[10,11]. It should be noted that we preferred this formula over the common Michaelis-Menten equation[1] as the range of $N_{int}$ in the biomass is limited, and the mathematical requirement of neglecting the half growth rate constant in high $N_{int}$ values ($K_N \ll N_{int,max}$) cannot be fulfilled.

$$f_{N_{int}} = \frac{\frac{N_{\text{int}} - N_{\text{int min}}}{N_{\text{int}}}}{\frac{N_{\text{int crit}} - N_{\text{int min}}}{N_{\text{int } crit}}} \text{ for } N_{int} < N_{\text{int crit}}, \text{ or } f_{N_{int}} = 1 \text{ for } N_{int} > N_{\text{int crit}} \quad \text{(S1.6)}$$

Where $N_{\text{int min}}$ and $N_{\text{int max}}$ (% g N g $DW^{-1}$) are the minimum and maximum internal nitrogen concentrations in *Ulva*, respectively, and $N_{crit}$ (% g N g $DW^{-1}$) is the threshold $N_{\text{int}}$ level below which the growth rate slows down.

*Nutrient Reserves.* Following the assumption of N limitation, internal N is the only modelled reserve. An underlying assumption is that the organic carbon reserve, depending on carbon uptake and photosynthesis rates, is not limiting within the modelled conditions (temperature ranges, seaweed density and water exchange and aeration rates). $N_{\text{int}}$ is affected by uptake of external N ($\psi_{N_{ext}}$, eq 2.1) and by dilution by growth ($N_{\text{int}}\mu$m).

$$\psi_{N_{ext}} = \frac{N_{\text{int max}} - N_{\text{int}}}{N_{\text{int max}} - N_{\text{int min}}} \frac{V_{\max} N_{\text{ext}}}{K_S + N_{\text{ext}}} \quad \text{(S2.1)}$$

Where $V_{\max}$ (µmol N g $DW^{-1}$ $h^{-1}$) is the maximum N uptake rate and $K_S$ (µmol N $l^{-1}$) is the N half saturation uptake constant.

*External Nutrients.* $N_{\text{ext}}$ represents the N concentration in the water in a cultivation reactor and is best described by the Convection–Diffusion equation[12] (eq S3.1).

$$\frac{\partial N_{ext}}{\partial t} = \nabla \cdot (D \nabla N_{ext}) - \nabla \cdot (\text{v} N_{ext}) - \psi_{N_{ext}} m \quad \text{(S3.1)}$$

Where $D$ ($m^2$ $s^{-1}$) is the average diffusivity coefficient of dissolved inorganic nitrogen species and v (m $s^{-1}$) is the velocity field in which the dissolved nitrogen is moving.

This equation can be simplified into eq S3, following a few assumptions: 1. $D$ is constant in space; 2. velocity flow is incompressible, and 3. net diffusivity is zero, as the reactor is well-mixed and there is no concentration gradient ($\nabla N_{ext} = 0$). Therefore, $N_{ext}$ in the reactor is affected only by the N supply by airlift pump (normalized to reactor volume) and N uptake by the algae.

*Model upscaling.* We upscaled the model in a two-step manner, as described in **Fig. S3**. On the first step, the metabolic model which applies to a single thallus scale (1 cm) is transitioned into a reactor scale (1 m) by adding an average light distribution term[8], allowing to adjust light intensity per mass to biomass density changes (eq S4). In this equation we multiplied $I_0$ by a 0.43 $PAR$ constant, representing the ratio of the sunlight which is suitable for photosynthesis[13]. Each reactor is assumed to be well-mixed by bottom aeration and is connected to an airlift pump, that supplies the reactor with fresh sea water and nutrients, as described by Chemodanov et al. [14]. We assume water flow through reactor boundaries is negligible.

$$f(I) = \frac{I_{average}}{K_I + I_{average}}, I_{average} = \frac{I_0 \, PAR}{K_0 Z + K_a SD} \left[1 - \exp\left(-(K_0 Z + K_a SD)\right)\right] \quad \text{(S4)}$$

Where $I_{average}$ (µmol photons $m^{-2}s^{-1}$) is the average photon irradiance in the reactor, $I_0$ (µmol photons $m^{-2}$ $s^{-1}$) is the incident photon irradiance at water surface, $SD$ (gDW $m^{-2}$) is the Stocking Density of biomass per unit of water surface in the reactor, $K_0$ ($m^{-1}$) is the water light extinction coefficient, Z (m) is the maximum water depth in the reactor, $K_a$ ($m^2$ $gDW^{-1}$) is the *Ulva* light extinction coefficient and $PAR$ is the Photosynthetically Active Radiation ratio of sunlight.

On the second step, the reactor scale model is transitioned into a farm scale model (1 km), simulating a row of reactors which are chained along the flow. Thus, each reactor constitutes a N sink, causing the spatial change of N concentrations in the environment. By assuming the width of this change is small in respect to the distance between the rows, a quasi-2D approximation can be made, making this model applicable also to multiple rows of reactors. This step requires the addition of a fourth governing equation, describing N dynamics in the environment, which is the connecting agent between the different reactors (eq S5). Each reactor is connected to the environment by the inflow of the airlift pump and the equal outflow of water surplus (eq S6), and the change in the environmental N concentrations depends on stream flow ($Q_s$) and environmental N concentrations ($N_{env}$), dilution ratio (*d*), pumping flow ($Q_P$) and uptake ($\psi_{N_{ext}}$) in reactors. All four ODEs were solved numerically with hourly time steps.

$$\frac{\partial N_{env}}{\partial t} = \frac{[-\mathbf{Q_s}(N_{env}(1-d)-N_{env})-\mathbf{Q_p}(N_{env}-N_{ext_x})]}{\boldsymbol{V_{cage}}} \quad \text{(S5)}$$

Initial and boundary conditions: $N_{env} = N_{env_{(x,t=0)}} = N_{env_0}$

$$\frac{\partial N_{ext}}{\partial t} = \frac{\mathbf{Q_p}(N_{env_x}-N_{ext_x})}{\boldsymbol{V_{cage}}} - \psi_{N_{ext}}\, m,\ \text{IC: } N_{ext_{(x,t=0)}} = N_{env_0} \quad \text{(S6)}$$

**Calculation of hourly growth and losses rates.** Maximum specific daily growth and losses rates were adjusted to hourly rates using eq S7. The formula relates to $\mu_{max}$ but can be used in the same way for $\lambda_{20}$. The equation was solved using the Microsoft Excel Office 365 solver.

$$m_0(1+\mu_{day,max}) = \sum_{i=0}^{\text{light hours}} m_i(1+\mu_{hour,max}) \quad \text{(S7)}$$

Where $m_0$ is the initial biomass density, $m_i$ is the biomass density in hour $i$, $\mu_{day,max}$ is the maximum specific daily growth rate, $\text{light hours}$ is the number of daily light hours and is set to 14 according to empiric data and $\mu_{hour,max}$ is the maximum specific hourly growth rate.

**Model calibration.** We calibrated the model parameters using experimental growth data of *Ulva* cultivation in a single well-mixed sea-based reactor from Chemodanov et al.[14] (**Figs S4-5 and Table S2**). First, we determined the *Ulva* light extinction coefficient, $K_a$, by minimizing the root mean relative error ($\text{RMSR}E_1$, eq S8) between modelled biomass growth from three experiments based on: 1. in situ measured light intensity (Onset HOBO Pendant®), and 2. light intensity data extracted from the IMS data base from the Israel Meteorological Services (https://ims.data.gov.il/he/ims/6). This was done by calculating $\text{RMSR}E_1$ for 20 values and 320 different parametric combinations of $\mu_{max}$, $K_a$, $K_I$, $n$, $T_{max}$, $T_{opt}$ and $\lambda_{20}$ in a defined range and identifying the $K_a$ value which results in the minimal errors**.** One experiment, where biomass degradation could not be explained by the model, was omitted from the calibration process. Next, using the same method, we determined the values of $\mu_{max}$, $K_I$, $n$, $T_{max}$, $T_{opt}$ and $\lambda_{20}$ (20 values and 280 different parametric combinations) by minimizing the mean error between measured and modelled biomass growth (N = 4) using in situ temperature data when available (in 3 out of 4 experiments) or IMS data otherwise, and IMS light intensity data ($\text{RMSR}E_2$, eq S9).

$$\text{RMSR}E_1 = \sqrt{\frac{\sum_{i=1}^{N}\left|\frac{PV_{m_{In}} - PV_{m_{Ex}}}{PV_{m_{In}}}\right|}{N}},\ N = 3 \quad \text{(S8)}$$

$$\text{RMSR}E_2 = \sqrt{\frac{\sum_{i=1}^{N}\left|\frac{m_f - PV_{m_{Ex}}}{m_f}\right|}{N}},\ N = 4 \quad \text{(S9)}$$

Where $\text{RMSR}E_1$ is the Root Mean Square Relative Error between $PV_{m_{In}}$ (g DW $l^{-1}$), the predicted value of final biomass based on in-situ light intensity data and $PV_{m_{Ex}}$ (g DW $l^{-1}$), the predicted value of final biomass based on ex-situ light intensity data, and $\text{RMSR}E_2$ is the Root Mean Square Relative Error between $m_f$ (g DW $l^{-1}$), the measured final biomass and $PV_{m_{Ex}}$.

**Model calibration data.** Data from returns 2, 4 and 5 was used for the calibration of the light extinction coefficient, $K_a$, and data from returns 1, 2, 4 and 5 was used for the calibration of $\mu$, $K_a$, $K_I$, $n$, $T_{max}$ and $T_{opt}$. Return 3 was not used for calibration as its negative growth could not be explained by the model.

**Model simulations.** *Simulated environmental conditions*. **Fig. S6** presents water temperature and light intensity profiles used for model simulations. Light intensity data was extracted from the IMS data base from the Israel Meteorological Services (https://ims.data.gov.il/he/ims/6). Ex-situ light intensity data was obtained at a 3-hour resolution and interpolated to a 1-hour resolution. Water temperature data was measured in-situ in a 1-month resolution and interpolated to a 1-hour resolution[15]. **Table S3** summarizes the main environmental data used for simulations and model boundary conditions. This data was collected and analyzed by Suari et al.[15] in a comprehensive study of the Alexander estuary. Salinity was set in a constant value of 30 PSU for two reasons. First, we found that in the examined range the model is not sensitive to the salinity level. Second, the study of Suari et al.[15] a complex estuary salinity dynamics, controlled by sandbar breaches rather than by seasonal phenomena, that does not allow describing salinity variations in an annual timeline manner.

### Results

#### Model calibration results.

*In- vs Ex-situ light intensity data and model projections.* In-situ light intensity and temperature data was obtained in a 15-minute resolution and averaged to a 1-hour resolution. Ex-situ light intensity data was obtained at a 3-hour resolution and interpolated to a 1-hour resolution. Temperature and light intensity profiles of returns 2, 4 and 5, followed by optimized models for each return based on both data sets, are presented in **Figs. S7-9.** Temperature and light intensity profile of returns 1, followed by an optimized model based on both ex-situ data is presented in **Fig. S10.** The in-situ based modelling uses eq S1.5 and the ex-situ based modelling uses eq S4.
*In- vs Ex-situ light intensity data parametric distribution.* $RMSRE_1$ was calculated for 320 parametric combinations of , $\mu_{max}$, $K_a$, $K_I$, $n$, $T_{max}$, $T_{opt}$ and $\lambda_{20}$ in a defined range (**Table S4**). Plotting $RMSRE_1$ vs the examined parameters (**Fig. S11**), points out $K_a$ as the only parameter with a clear trend, achieving minimal errors around $K_a = 0.15$.
Next, after setting $K_a$ as 0.15, $RMSRE_2$ was calculated for 280 parametric combinations of $\mu_{max}$, $K_I$, $n$, $T_{max}$, $T_{opt}$ and $\lambda_{20}$ in the same defined range (**Table S4**). Plotting $RMSRE_2$ vs these 320 examined parametric combinations (**Fig. S12**) suggests setting $\lambda_{20}$ and $K_I$ on the low side of the examined range. Therefore, and after manual examination of specific parametric combinations, we set $K_I = 20$ and $\lambda_{20} = 0.0016$.

*Model parameters.* Details regarding model parameters, including original values, examined ranges and chosen values, are presented in **Table S4**. **Table S5** presents the detailed calibration results, comparing measured and modeled biomass production in returns 1,2,4 and 5, and average and specific relative errors.

**Model simulations.** *Seasonal trends in biomass production and nitrogen removal.* **Fig. S13** presents how shortening the cultivation cycles during spring and autumn can enable maintaining a constant chemical composition year-round.

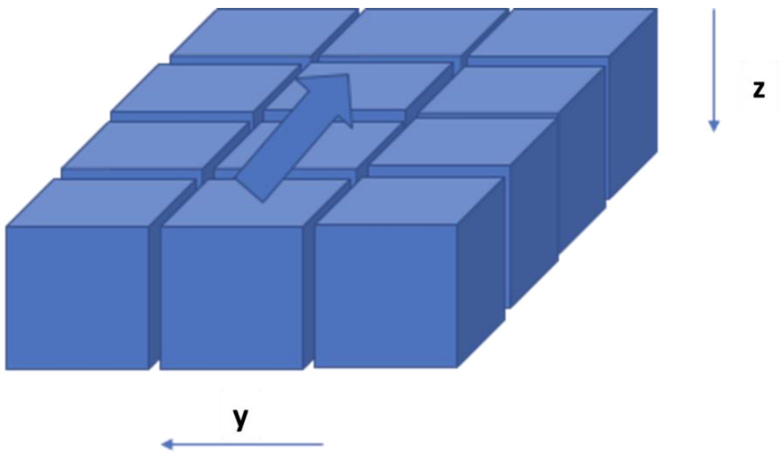


**Fig. S1.** Dimensions of the multi-scale model: x axis is the direction of the flow, along which cultivation reactors are chained, y axis is the horizontal direction perpendicular to the flow, along which multiple rows of reactors can be placed, and z axis points downwards from the water surface. Only one layer of reactors can be placed along the z axis, as sun exposure is necessary.

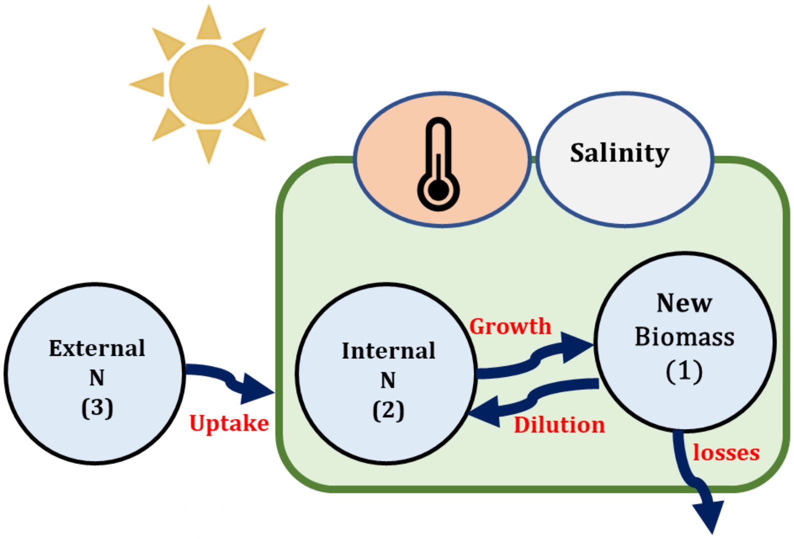


**Fig. S2.** A schematic description of the metabolic model of Ulva. The rounded rectangle represents the algae. The surrounding area represents the environment to which the algae is exposed. Circles represent the model state variables and constraining environmental conditions are represented by ellipses and icons: light intensity with a direct effect on growth rate and salinity and temperature that affect all biological processes.

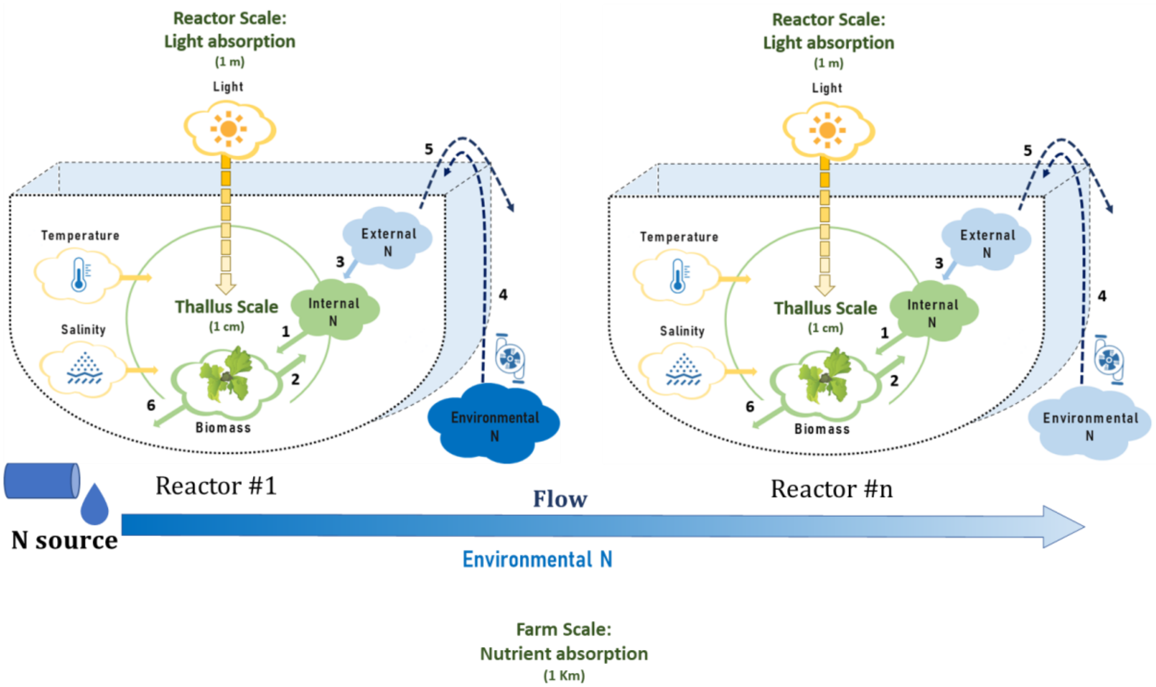


**Fig. S3.** A schematic description of the multi-scale model. The thallus scale (1 cm, green circle) is composed of a simple metabolic model of Ulva, in which the production of new biomass (Ulva icon) is affected by internal nitrogen (N, full green cloud) and by constraining environmental conditions, including light intensity, salinity and temperature (yellow clouds). The reactor scale (1 m, U shape pictures) adds light extinction effects (yellow graduated arrow), the concentration of external N in the reactor and the concentration of environmental N outside the reactor (dark/light blue clouds, depending on N concentration). The farm-scale (1 km, row of reactors starting at Reactor #1 and counting downstream to Reactor #n) adds the nutrient reduction caused by absorption in reactors along with the flow (Blue graduated arrow). Green and blue clouds represent the model state variables. Numbers represent the following processes: 1. Biomass growth; 2. Dilution of internal N by growth; 3. N uptake; 4-5. Water exchange by airlift pumping and overflow, and 6. Biomass losses.

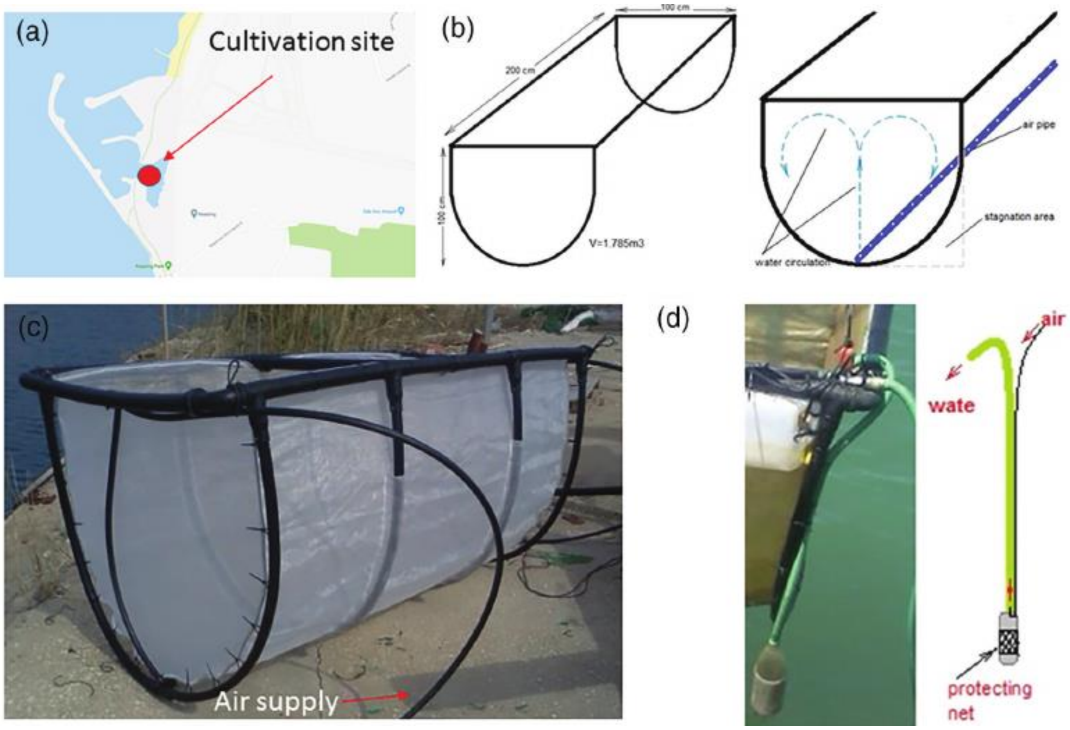


**Fig. S4.** (a) Cultivation site; (b) schematic design of the reactor with intensification with tumbling, mixing and water exchange; (c) image of the reactor for intensified cultivation, and (d) external airlifts for water exchange enhancement. (Adapted from Chemodanov et al. [14] with permission).

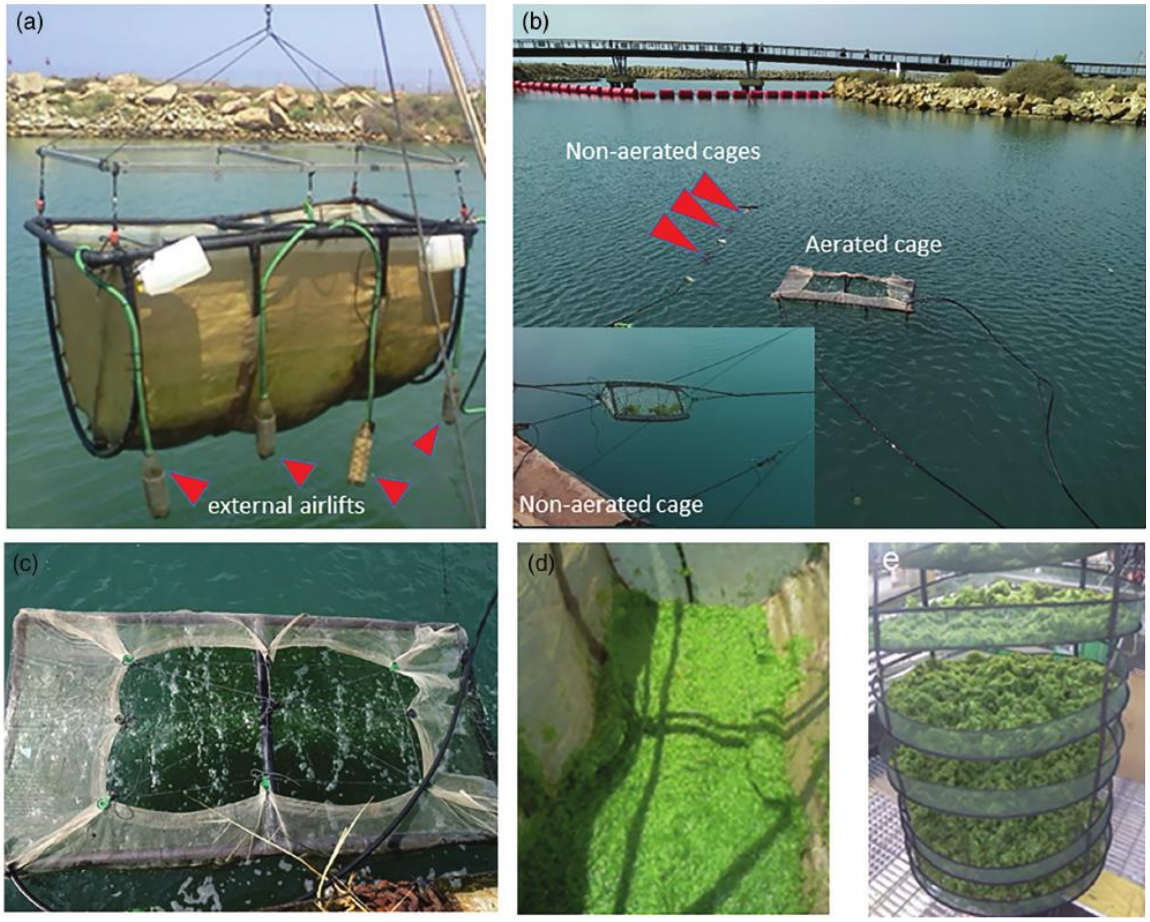


**Fig. S5.** (a) Image of the cultivation reactor with external airlifts; (b) deployment of the reactor with algae to the cultivation site; (c) tumbling with air and mixing of *ulva* sp. biomass in the reactor; (d) harvested *Ulva* biomass after water removal with gravitation and (e) solar dried *Ulva* biomass (Adapted from Chemodanov et al. [14] with permission).

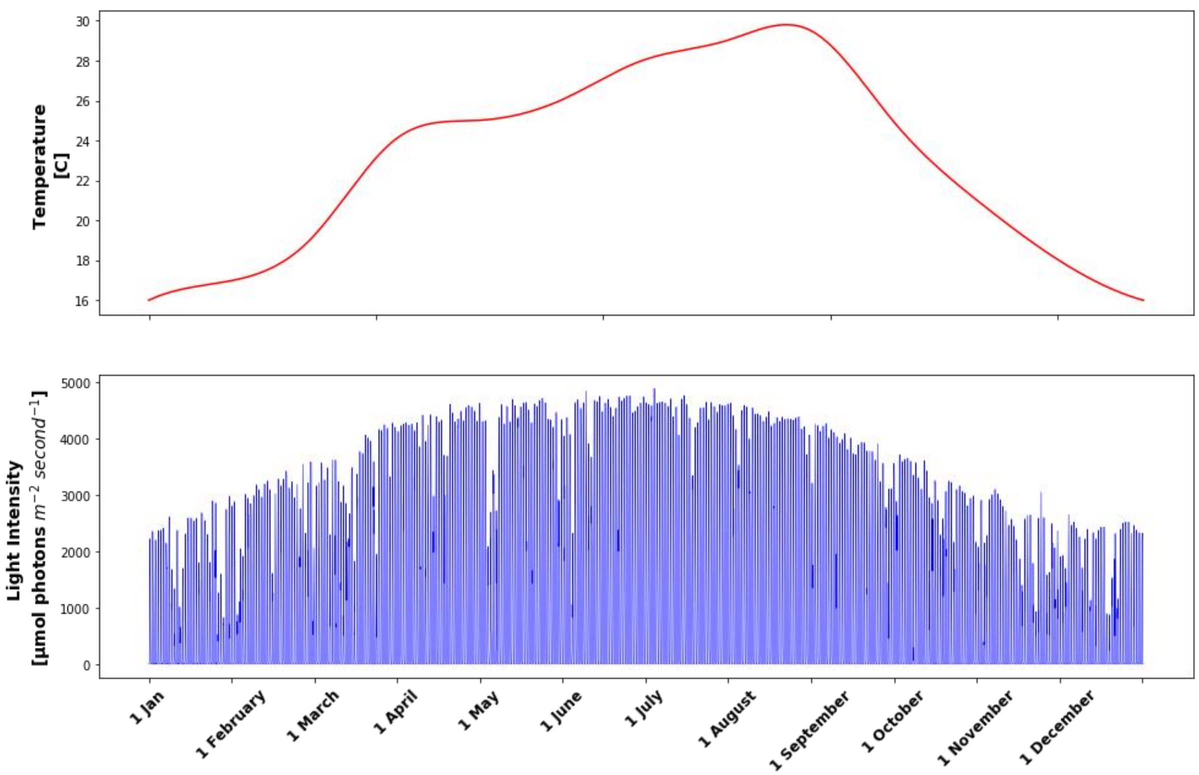


**Figure S6.** Water temperature (red) and light intensity (blue) profiles of the simulated environment.

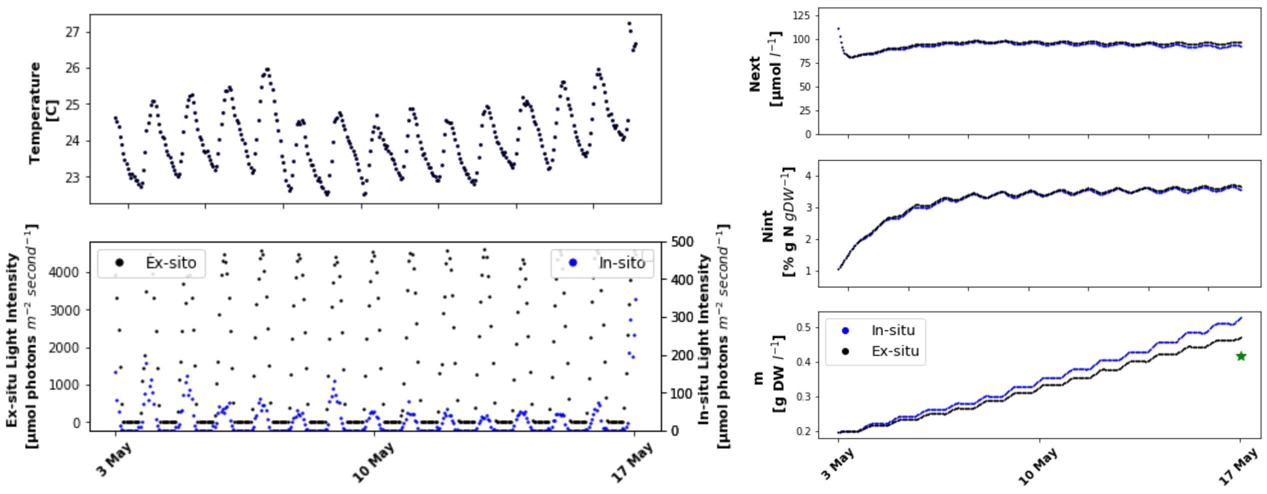


**Fig. S7.** Return 2. left: Temperature and light intensity profile. Right: Model results based on in-situ (blue) and ex-situ (black) light intensity measurements. Green star marks measured final m.

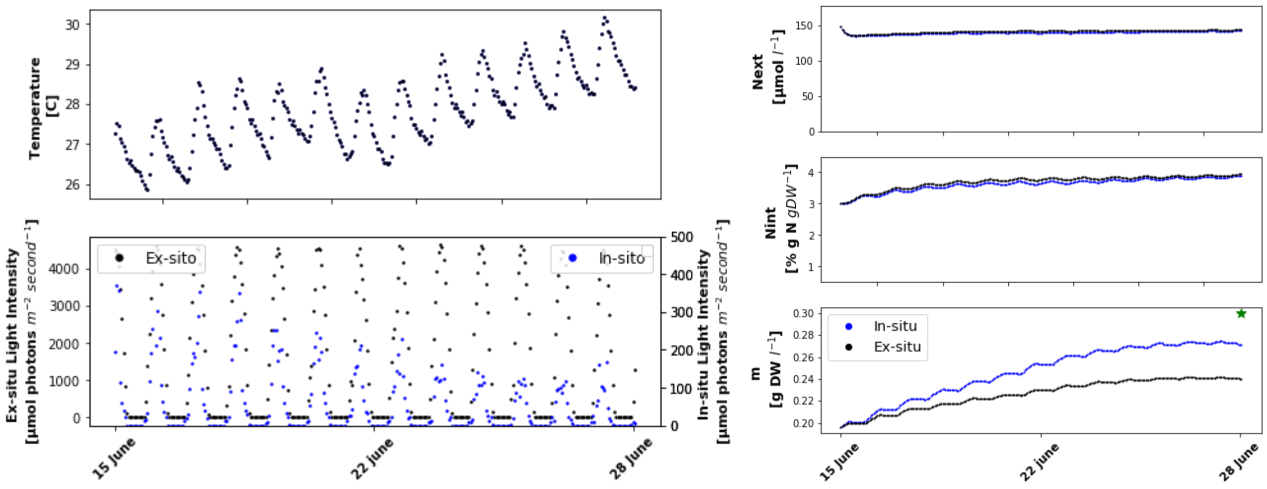


**Fig. S8.** Return 4. left: Temperature and light intensity profile. Right: Model results based on in-situ (blue) and ex-situ (black) light intensity measurements. Green star marks measured final m.

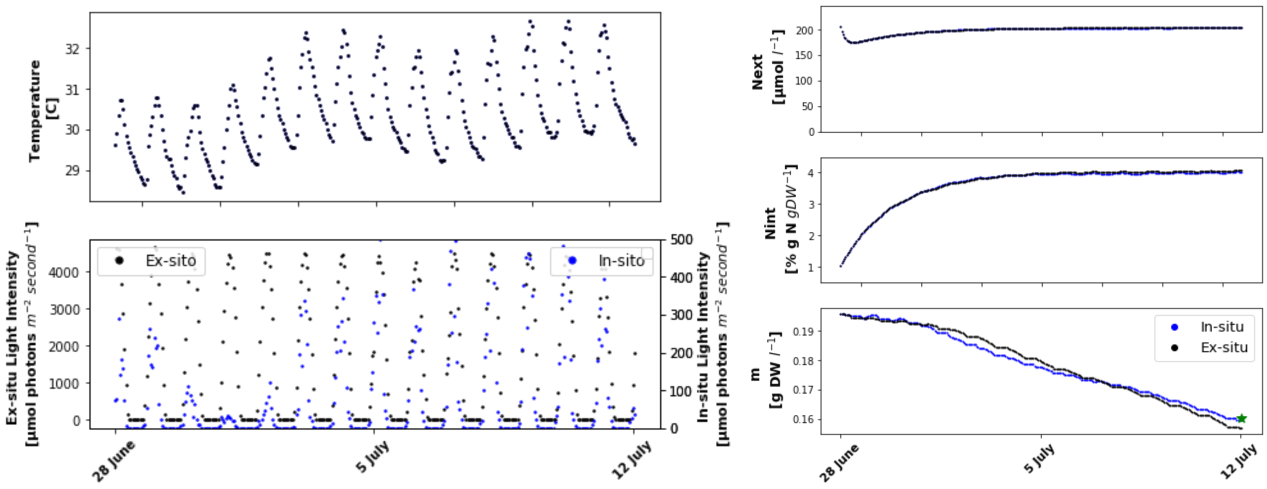


**Fig. S9.** Return 5. left: Temperature and light intensity profile. Right: Model results based on in-situ (blue) and ex-situ (black) light intensity measurements. Green star marks measured final m.

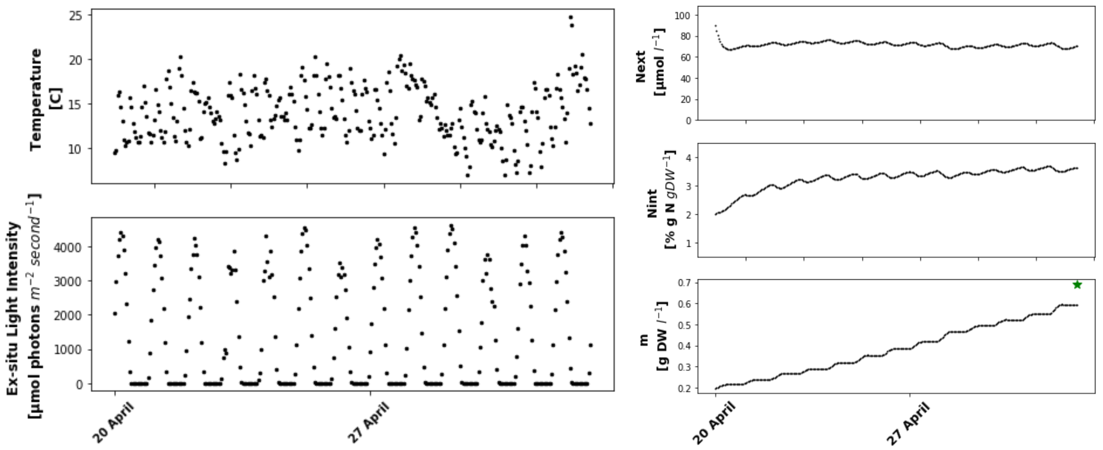


**Fig. S10.** Return 1. left: Temperature and light intensity profile based on ex-situ measurements. Right: Model results based on ex-situ temperature and light intensity measurements. Green star marks measured final m.

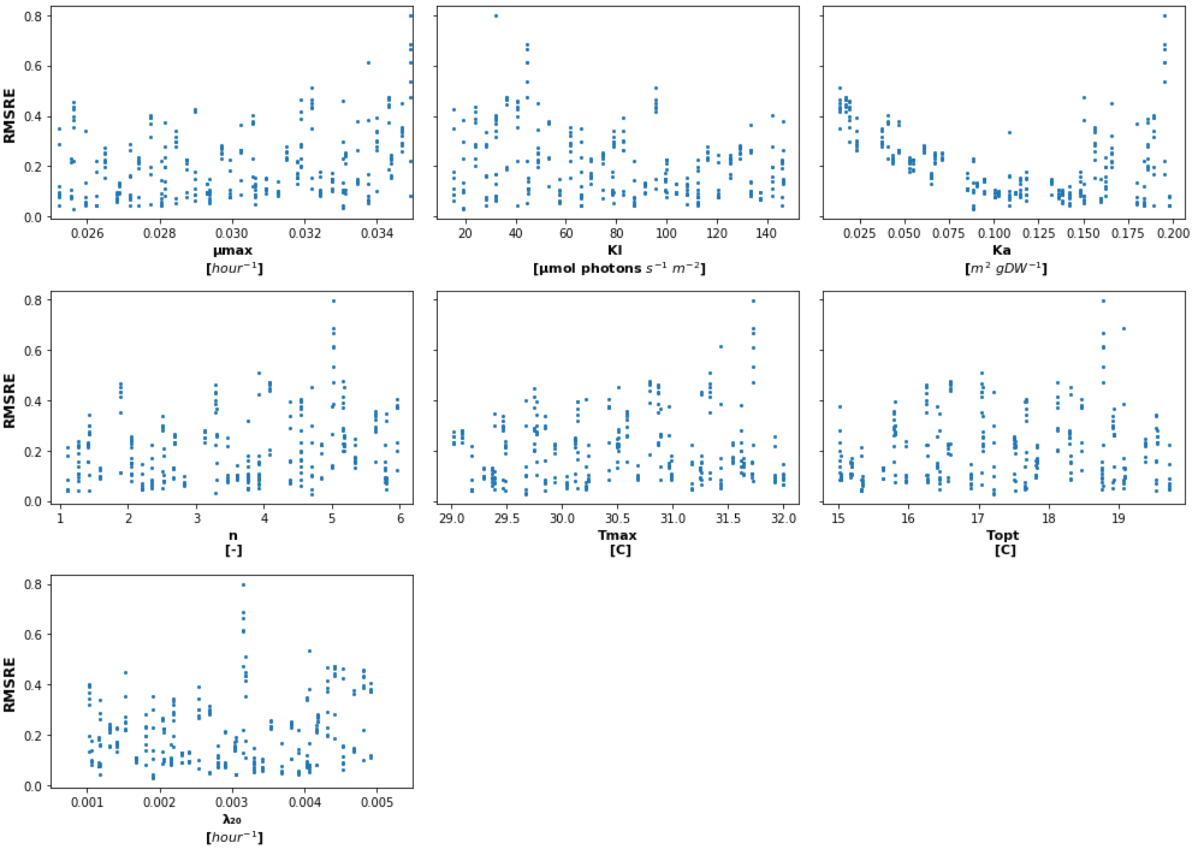


**Fig. S11.** $RMSRE_1$ distribution for parametric combinations of μ, $\lambda_{20}$, $K_a$, $K_I$, $n$, $T_{max}$ and $T_{opt}$ in a defined range (See **Table S2**).

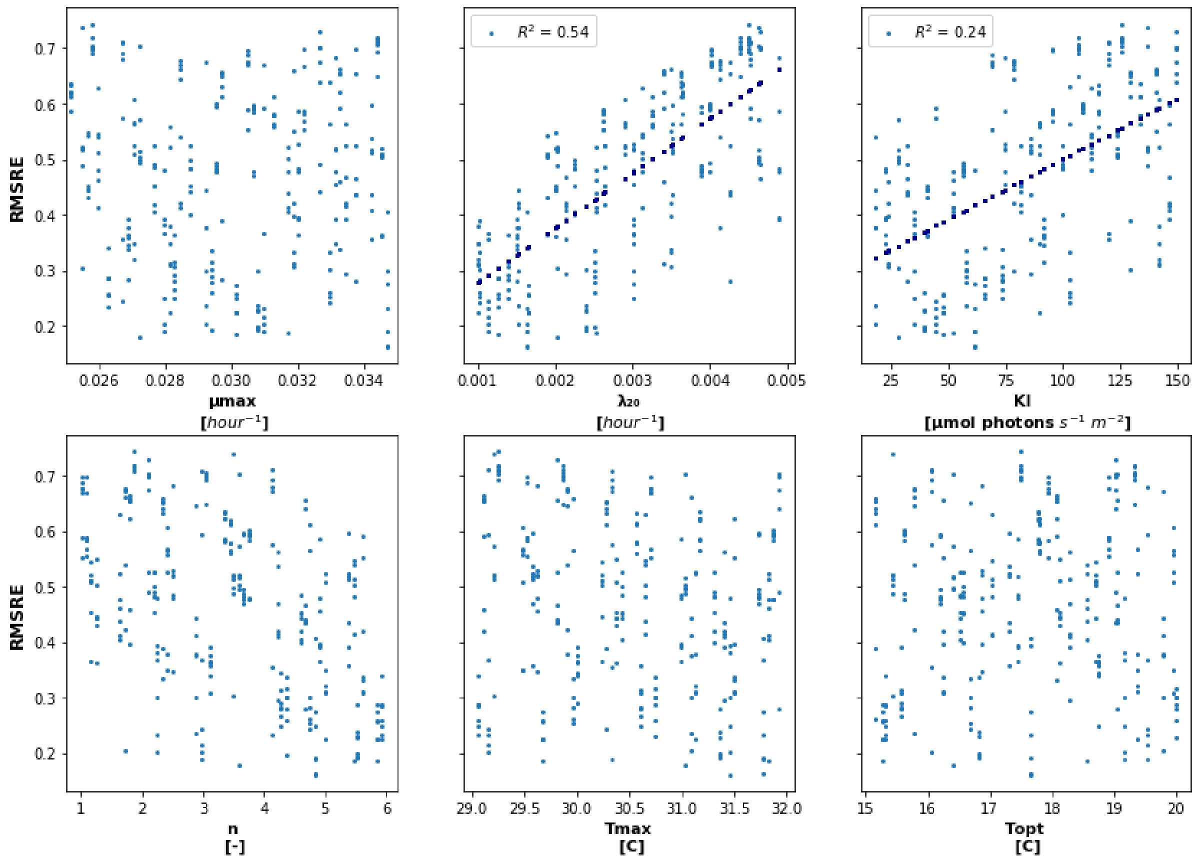


**Fig. S12.** $RMSRE_2$ distribution for parametric combinations of μ, $\lambda_{20}$ , $K_I$, $n$, $T_{max}$ and $T_{opt}$ in a defined range (See **Table S2**).

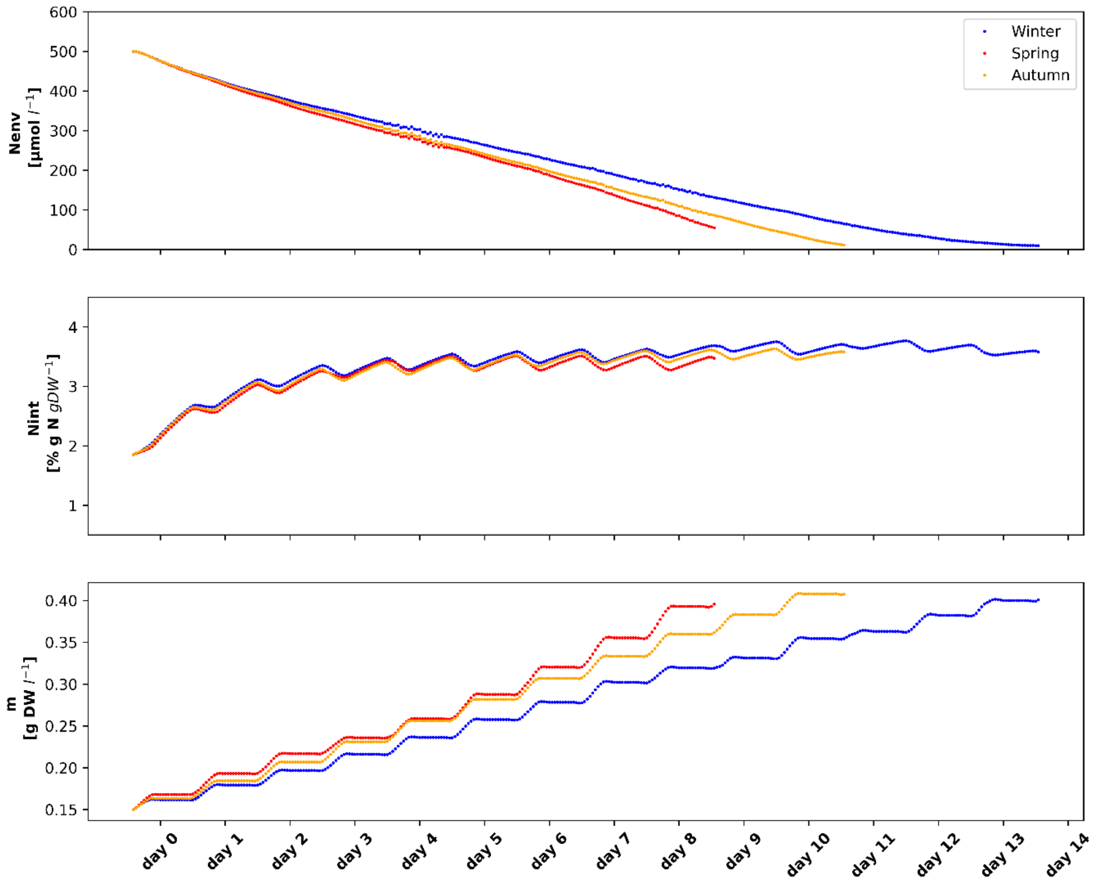


**Fig. S13**. $N_{\mathrm{env}}$, $N_{\mathrm{int}}$ and $m$ dynamics along seasonal specific cultivation periods: 14 days in winter, 9 days in spring and 11 days in autumn, for the last reactor in a farm of 731 cages

**Table S1.** Metabolic model governing equations

| State Variable | ODE | Equation | Annotation |
|---|---|---|---|
| $m$ | $\frac{\partial m}{\partial t} = (\mu - \lambda)\, m$<br>Initial Condition (IC):<br>$m_{(t=0)} = m_0$ | (eq S1) | μ is biomass specific growth rate as a function of I, T, S and nutrients [1] (eq S1.1 and S1.3-S1.6) and $\lambda$ is biomass specific losses rate as a function of T (eq S1.2) |
| $N_{\mathrm{int}}$ | $\frac{\partial N_{int}}{\partial t} = \psi_{N_{ext}} - N_{\mathrm{int}}\mu\mathrm{m}$<br>IC: $N_{\mathrm{int}\ (t=0)} = N_{\mathrm{int}_0}$ | (eq S2) | $\psi_{N_{ext}}$ is the N uptake function (eq S2.1) and $N_{\mathrm{int}}\mu\mathrm{m}$ represents dilution of internal N by growth |
| $N_{\mathrm{ext}}$ | $\frac{\partial N_{ext}}{\partial t} = -\frac{\nabla\cdot(\mathbf{v}N_{ext})}{V_{cage}} - \psi_{N_{ext}} m$<br>IC: $N_{\mathrm{ext}\ (t=0)} = N_{\mathrm{ext}_0}$ | (eq S3) | $-\frac{\nabla\cdot(\mathbf{v}N_{ext})}{V_{cage}}$ represents the convection of $N_{ext}$ into or out of the reactor and $\psi_{N_{ext}} m$ represents the N sink in the biomass. Eq S3 derives from the Convection-Diffusion equation [12] as described below (eq S3.1). |

**Table S2.** Experimental data used for model calibration [14]

| Return | Starting date (2017) | Final date (2017) | Initial biomass* [g FW m$^{-3}$] | Final biomass* [g FW m$^{-3}$] | Initial $N_{\mathrm{int}}$ [% g N g DW$^{-1}$] | Final $N_{\mathrm{int}}$ [% g N g DW$^{-1}$] | $N_{ext}$ [µmol N l$^{-1}$] | Light intensity data source |
|---|---|---|---|---|---|---|---|---|
| 1 | 20 April | 3 May | 1,315 | 4,590 | N/A, 2** | 1.05 | 90 | Ex-situ |
| 2 | 3 May | 17 May | | 2,781 | 1.05 | 0.65 | 111 | In + Ex-situ |
| 3 | 17 May | 29 May | | 1,120 | 0.65 | 0.98 | N/A, 91*** | In + Ex-situ |
| 4 | 15 June | 29 May | 1,306 | 2,001 | N/A, 3** | 1.06 | 148 | In + Ex-situ |
| 5 | 28 June | 12 July | | 1,069 | 1.06 | 1.40 | 206 | In + Ex-situ |

*DW:FW conversion ratio of 0.15

**Assumed based on personal communication (A. Chemodanov, April 2020)

*** Used final $N_{\mathrm{ext}}$ values instead of N/A initial values

**Table S3.** Simulated environmental conditions. Based on data from the Alexander estuary [15]

| Parameter | Unit | Value |
|---|---|---|
| Dissolved N | μmol N $l^{-1}$ | 500 |
| Dissolved P | μmol P $l^{-1}$ | 50 |
| Bottom Salinity | PSU | 12-30* |
| Stream flow | l $hour^{-1}$ | 7776** |
| Estuary width | m | 22 |
| Estuary depth | m | 2.5 |

*Used a constant salinity value of 30 PSU

**Calculated based on a baseflow controlled by a stable influx of 11.5 $10^3$ $m^3$ $day^{-1}$, normalized to a per hour flow through an area equal to the narrow-side cage cross section (0.89 $m^2$), in defined estuary width and depth.

**Table S4.** Model parameters: literature values, examined ranges and chosen values

| Parameter | Literature values | Unit | Reference | Examined range | Best value | unit | Determining Analysis |
|---|---|---|---|---|---|---|---|
| $\mu_{max}$ | 0.416 (0.027) | $day^{-1}$ (light $h^{-1}$) | [8] | 0.025-0.035 | * | Light $h^{-1}$ | |
| $\lambda_{20}$ | 0.066 (0.0049) | $day^{-1}$ (light $h^{-1}$) | [8] | 0.014-0.068 (0.001-0.005) | 0.022 (0.0016) | $day^{-1}$ (Light $h^{-1}$) | 2 |
| $\theta$ | 1.047 | unitless | [1] | 0.9-1.2 | * | | |
| $N_{\text{int max}}$ | 4.2 | % Kg N Kg $DW^{-1}$ | ** | 3.2-4.5 | * | | |
| $N_{\text{int min}}$ | 0.7 | % Kg N Kg $DW^{-1}$ | [1] | 0.5-0.7 | * | | |
| $N_{\text{int } crit}$ | 2 | % Kg N Kg $DW^{-1}$ | [16] | 1.5-3 | * | | |
| $V_{max}$ | 60 | μmol N g $DW^{-1}$ $h^{-1}$ | [17–19] | 50-250 | * | | |
| $K_S$ | 14 | mmol N $m^{-3}$ | [18,20] | 10-30 | * | | |
| $K_I$ | 144.8 | μmol photons $m^{-2}$ $s^{-1}$ | [8] | 15-150 | 20 | μmol photons $m^{-2}$ $s^{-1}$ | 2 |
| $K_0$ | 1.5 | $m^{-1}$ | [8] | 0.1-3 | * | | |
| $K_a$ | 0.01 | $m^2$ $gDW^{-1}$ | [8] | 0.01-0.2 | 0.15 | $m^2$ $gDW^{-1}$ | 1 |
| $T_{min}$ | 5 | ℃ | [1] | 1-10 | * | | |
| $T_{opt}$ | 25 | ℃ | [1] | 15-20 | 18 | ℃ | |
| $T_{max}$ | 35 | ℃ | [1] | 29-32 | 31.5 | ℃ | |
| n | 2 | - | [1] | 1-6 | 2 | | 2 |
| $S_{min}$ | 0 | PSU | [1] | 0-10 | * | | |
| $S_{opt}$ | 18 | PSU | [1] | 15-25 | * | | |
| $S_{max}$ | 45 | PSU | [1] | 40-50 | * | | |
| $Q_P$ | 460 | l $h^{-1}$ | [14] | 300-600 | * | | |
| $d$ | 0 | % $reactor^{-1}$ | | 0-5 | * | | |

*kept original literature/assumed value unchanged

**$N_{\text{int max}}$ value determined based on personal observations

**Table S5.** Calibration results: measured and modeled biomass production in returns 1,2,4 and 5

| Return | Initial biomass [g FW $m^{-3}$] | Measured Final biomass [g FW $m^{-3}$] | Modeled Final biomass: In-Situ data [g FW $m^{-3}$] | Modeled Final biomass: Ex-Situ data [g FW $m^{-3}$] | Relative Error 1* | Relative Error 2** |
|---|---|---|---|---|---|---|
| 1 | 1,315 | 4,590 | | 3,961 | | 0.137 |
| 2 | | 2,781 | 3,518 | 3,137 | 0.121 | 0.128 |
| 4 | 1,306 | 2,001 | 1,810 | 1,602 | 0.129 | 0.200 |
| 5 | | 1,069 | 1,064 | 1,048 | 0.017 | 0.021 |
| RMSRE | | | | | 0.103 | 0.138 |

*Calculated by eq S8

**Calculated by eq S9
